# Effects of time-dependent ATP consumption caused by neuron firing on ATP concentrations in synaptic boutons containing and lacking a stationary mitochondrion

**DOI:** 10.1101/2024.02.20.581271

**Authors:** Andrey V. Kuznetsov

## Abstract

The precise mechanism behind the absence of a stationary mitochondrion in approximately half of presynaptic release sites in axons, and how these sites lacking a stationary mitochondrion receive ATP, is not fully understood. This paper presents a mathematical model designed to simulate the transient ATP concentration in presynaptic *en passant* boutons. The model is utilized to investigate how the ATP concentration responds to increased ATP demand during neuronal firing in boutons with a stationary mitochondrion and those without one. The analysis suggests that neuron firing may cause oscillations in the ATP concentrations, with peak-to-peak amplitudes ranging from 0.06% to 5% of their average values. However, this does not deplete boutons lacking a mitochondrion of ATP; for physiologically relevant values of model parameters, their concentration remains approximately 3.75 times higher than the minimum concentration required for synaptic activity. The variance in average ATP concentrations between boutons containing a stationary mitochondrion and those lacking one ranges from 0.3% to 0.8%, contingent on the distance between the boutons. The model indicates that diffusion-driven ATP transport is rapid enough to adequately supply ATP molecules to boutons lacking a stationary mitochondrion.

## 1. Introduction

The primary role of mitochondria is to produce easily accessible chemical energy in the form of ATP [1,2]. This occurs because mitochondrial respiration generates a substantially larger amount of ATP from a glucose molecule compared to glycolysis, with a ratio of 36:2.

Apart from stationary mitochondria, some mitochondria are transported in anterograde and retrograde directions [3-6]. These mobile mitochondria contribute to supplying ATP to energy-demanding sites, such as *en passant* boutons (hereafter referred to as boutons). However, it is believed that the primary source of ATP for these boutons comes from stationary mitochondria that specifically dock in them [7-9].

In the brain, synapses represent the main locations where ATP is consumed [10]. The reason why approximately half of the presynaptic release sites contain a stationary mitochondrion remains a mystery, while the remaining sites lack one [11-14]. The question of how a bouton without a stationary mitochondrion obtains its supply of ATP is investigated in this paper.

Three competing hypotheses in the literature propose explanations for this phenomenon: ATP is transported to boutons lacking mitochondria through diffusion [15], ATP is synthesized in boutons lacking mitochondria via glycolysis [15], and ATP needs are partially fulfilled by ATP production in neighboring axons, dendrites, and glial cells [12,16].

Choosing among these three models requires investigating whether diffusion-driven ATP transport alone is sufficiently fast to support ATP demand in a bouton lacking a mitochondrion. If diffusion is fast enough, the ATP concentration will remain relatively uniform; if it is slow, the ATP concentration will decrease with distance from a mitochondrion and fall below the threshold needed to sustain synaptic activity in a bouton lacking a mitochondrion [12,16].

Compartmental models are frequently employed in synapse modeling. Ref. [17] developed a model comprising 33 ordinary differential equations, which simulates neuronal, astrocytic, extracellular, and vascular compartments. In ref. [18], a neurochemical model of a tripartite synapse, consisting of a presynaptic terminal, post-synaptic terminal, and astrocyte, was developed, which involves 143 ordinary differential equations. This model predicts the occurrence of local sleep, which is crucial for cellular resource restoration.

In contrast to previous models, the model developed here not only tracks ATP variation over time but also over position in the axon. This enables the simulation of time-dependent differences in ATP concentration between synapses with and without a stationary mitochondrion, located at different positions along the axon. In contrast to our previous work [19], which solely simulated a steady-state scenario, this study simulates a scenario where ATP consumption fluctuates due to neuron firing.

The frequency of neuron firing usually falls within the range of 1 to 200 Hz [20]. The rate of ATP production by mitochondria fluctuates over the course of the day [21]. However, mitochondria are unlikely to adjust ATP production at a pace equivalent to neuron firing. According to ref. [22], the estimated response time for mitochondrial oxygen consumption to stepwise changes in cardiac energy demand is 4-8 seconds. Ref. [23] suggests that it takes minutes to change the ATP production rate in mitochondria. The significant reliance on glycolysis for ATP production during high-intensity exercise, as noted in ref. [24], implies that ATP production by mitochondria cannot be increased within seconds. Considering that ATP cannot be stored and must be generated as needed [25], it is important to investigate the extent of ATP concentration variations in boutons due to changes in ATP demand during neuron firing.

## 2. Materials and models

### 2.1. Equations governing ATP concentration in a single periodic unit control volume (CV) containing a bouton having a stationary mitochondrion and a bouton lacking a stationary mitochondrion

Stationary mitochondria are uniformly distributed along the length of an axon [26]. The model postulated that a stationary mitochondrion is present at the center of every other varicosity (bouton) along the axon. This simplifying assumption is corroborated by two experimental observations described above: (i) approximately half of the presynaptic release sites contain a stationary mitochondrion, and (ii) the distribution of stationary mitochondria is uniform. The assumption of periodicity enabled the definition of a periodic unit CV containing half of a mitochondrion on each side (the other half belonging to a neighboring periodic CV), as illustrated in Fig. 1. The width of axonal varicosity is 2*δ*, while the average spacing between varicosities is *L*. Stationary mitochondria are assumed to be present at every second varicosity, with a distance of 2 *L* + 4*δ* between them.

**Fig. 1.**
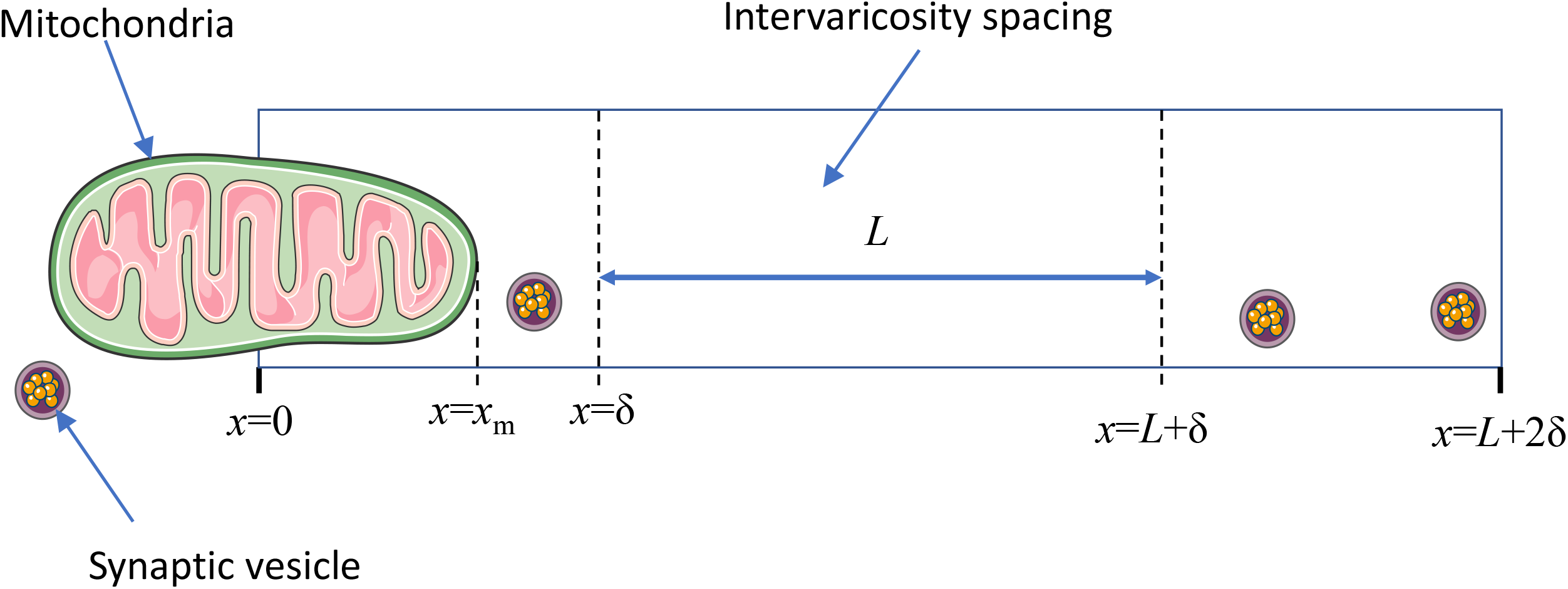
(a) A schematic diagram illustrates half of a periodic unit CV. The displayed segment contains half of a mitochondrion, with the other half belonging to a neighboring periodic CV. The width of axonal varicosity is denoted by 2*δ* ; the average spacing between varicosities is denoted by *L*. It is assumed that stationary mitochondria are positioned at every other varicosity. The spacing between stationary mitochondria is 2 *L* + 4*δ*. *Figure generated with the aid of servier medical art, licensed under a creative commons attribution 3.0 generic license*. http://Smart.servier.com.

Due to symmetry, only half of the CV, displayed in Fig. 1, needs to be analyzed. In the segment of the varicosity where a stationary mitochondrion is located (Fig. 1), the conservation of ATP molecules leads to the following equation:

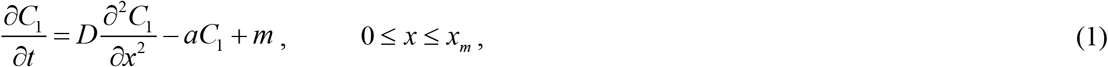

where *t* denotes the time and *x* denotes the linear coordinate along the axon. Eq. (1) assumes that ATP consumption is proportional to the amount of ATP available for binding.

In the segment of varicosity not occupied by a stationary mitochondrion (Fig. 1), the conservation equation for ATP molecules is as follows:

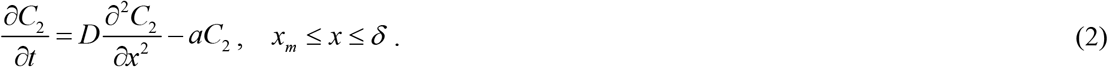

In the space between varicosities, known as intervaricosity spacing (Fig. 1), the conservation of ATP leads to the following equation:

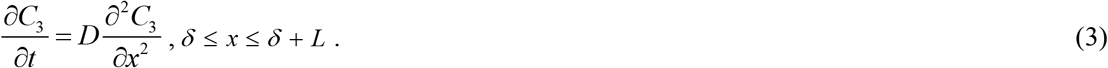

In an empty varicosity, where ATP is still consumed but lacks a stationary mitochondrion, resulting in no ATP production (Fig. 1), the conservation of ATP leads to the following equation, similar to Eq. (2):

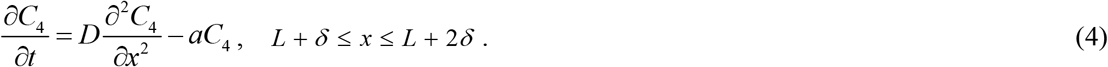

The only dependent variable in the model is the linear concentration of ATP in the axon, denoted as *C*. The model parameters are summarized in Table 1.

**Table 1.** Parameters describing ATP transport and accumulation in the axon.

| Symbol | Definition | Unit | Value or range | Reference(s) | Value used in computations (unless a different value is indicated in the figure caption) |
| --- | --- | --- | --- | --- | --- |
| $a_0$ | Kinetic constant describing the rate of ATP consumption in a bouton | $s^{-1}$ | 0.23 <sup>a</sup> | | 0.23 |
| $A_c$ | Cross-sectional area of an axon | $\mu\text{m}^2$ | 0.79 <sup>b</sup> | | 0.79 |
| $C_0$ | Linear ATP concentration in the axon | $\mu\text{m}^{-1}$ | $1.42 \times 10^6$ <sup>c</sup> | [15] | $1.42 \times 10^6$ |
| $\hat{C}_0$ | Volumetric ATP concentration in the axon | mM | 2-4 | [15] | 3 |
| $C_{\min}$ | Minimum linear ATP concentration needed to support synaptic transmission | $\mu\text{m}^{-1}$ | $3.78 \times 10^5$ <sup>d</sup> | [15] | $3.78 \times 10^5$ |
| $D$ | Diffusivity of ATP in the cytosol | $\mu\text{m}^2 \text{ s}^{-1}$ | 710-720<br>in water,<br>~300 in the cytosol | [29,30] | 300 |
| $f$ | Frequency at which neuron fires | Hz | 1-200 | [20] | 1, 10 |
| $L$ | Average spacing between varicosities | $\mu\text{m}$ | 3 | [12] | 3, 10 |
| $m$ | Rate of ATP production per unit length of a mitochondrion | $\mu\text{m}^{-1} \text{ s}^{-1}$ | $1.29 \times 10^6$ <sup>e</sup> | [31-33] | $1.29 \times 10^6$ |
| $\dot{m}$ | Rate of ATP production per unit mass of tissue | mol/(s kg) | $1.1 \times 10^{-4}$<br>$-1.83 \times 10^{-4}$ <sup>f</sup> | [31] | $1.33 \times 10^{-4}$ |
| $2x_m$ | Length of a mitochondrion | $\mu\text{m}$ | 0.5-10 | Fig. 2G of [12], [34] | 1 |
| $\gamma$ | Percent of mitochondria by volume in tissues | % | 2-10 | [33] | 5 |
| $\delta$ | Half-width of axonal varicosity | $\mu\text{m}$ | 1 | Fig. 2E of [12], [35] | 1 |
| $\varpi$ | Mass of a single mitochondrion | kg | $322 \times 10^{-15}$ | [32] | $322 \times 10^{-15}$ |

<sup>a</sup> The value of $a_0$ was found by using the analytical solution given in section S1 of Supplementary Materials. The condition used was that the maximum value of ATP concentration at a constant rate of ATP consumption in boutons, occurring at $x = 0$ , is equal to $C_0$ .
<sup>b</sup> The estimation of $A_c$ proceeded as follows:
$A_c = \frac{\pi d^2}{4}$ . A diameter of 1 $\mu\text{m}$ was used for the axon diameter, $d$ . Cross-sectional area $A_c$ was assumed to be constant in our model.
<sup>c</sup> This value was estimated as follows:
<sup>d</sup> To sustain synaptic transmission, boutons must sustain a minimum ATP concentration of around 0.8 mM [15]. This corresponds to
<sup>e</sup> The quantity of ATP molecules produced by mitochondria per unit length of the axon is approximated as:
<sup>f</sup> The range for $\dot{m}$ given in ref. [31] is between $6.6 \times 10^{-3}$ - $11 \times 10^{-3}$ mol/(min kg).

Given that a significant portion (roughly 80%) of energy in mammalian cortex neurons is expended on action potentials [27], it is anticipated that ATP consumption will fluctuate over time. To simulate the fluctuating ATP consumption rate required to sustain synaptic activity during neuronal firing, the following periodic ATP consumption rate is assumed:

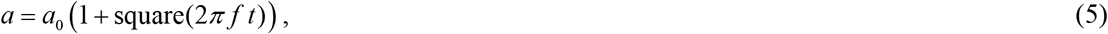

where square(2*π ft*) is a periodic step function with a period of 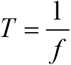. It resembles the sine function but produces a square wave with values of –1 and 1.

Fig. 2 illustrates the function 1 + square(2*π f t*) for *f* = 1 Hz during the initial 10 seconds. Eq. (5) follows an all-or-nothing approximation. A basal level of ATP consumption is necessary in a synapse for homeostatic maintenance [28]. However, as the objective of this paper is to estimate the maximum amplitude of ATP concentration oscillations in boutons with and without a stationary mitochondrion in response to neuron firing, the approximation given by Eq. (5) is justified.

**Fig. 2.**
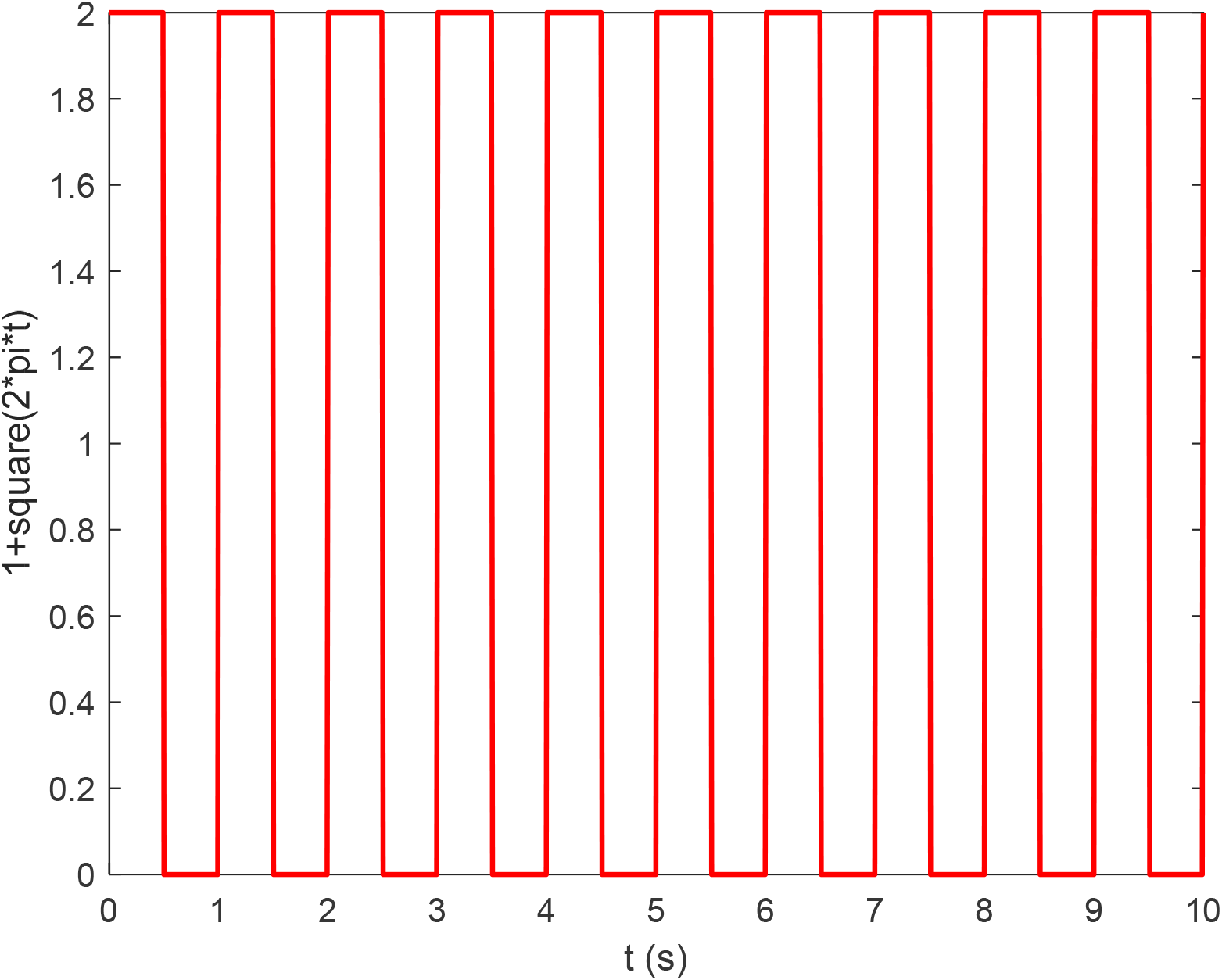
A step function of time is employed to simulate the oscillation in the rate of ATP consumption in a bouton resulting from neuron firing, as described in Eq. (5), *f* =1 Hz. Only the first 10 seconds are depicted; thereafter, the pattern continues in the same manner until the end of the simulation (*t* _*f*_ = 1000 s). When considering constant ATP consumption, this oscillating step function is replaced with unity, resulting in *a* = *a*_0_.

Eqs. (1)-(4) were solved with the following boundary conditions:

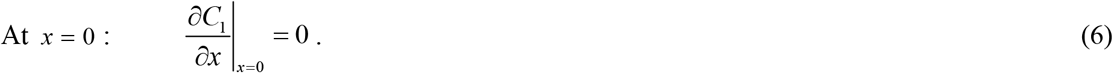

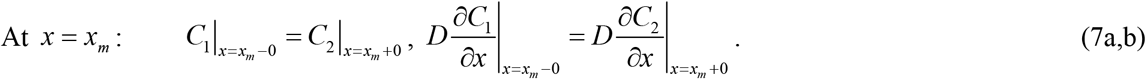

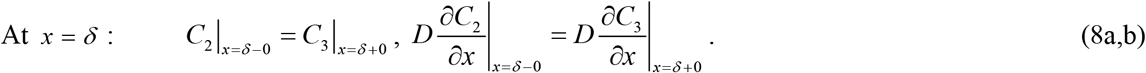

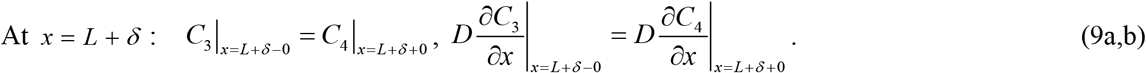

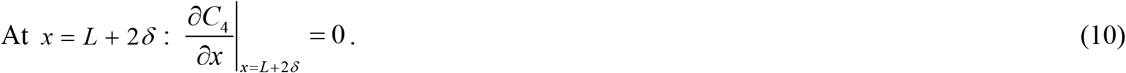

ATP diffusivity is incorporated into the left- and right-hand sides of Eqs. (7b), (8b), and (9b) to allow for potential model extensions in the future. This allows for the possibility of different diffusivity values in various subdomains.

Zero initial conditions were employed:

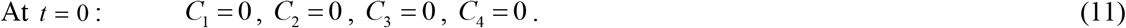

### 2.2. Numerical solution

Eqs. (1)-(4) constitute a system of partial differential equations (PDEs) that can be solved by a single-domain approach [36]. This method employs a single PDE with coefficients that change at subdomain interfaces. As a result, the equation adjusts to the appropriate form within each of the four subdomains:

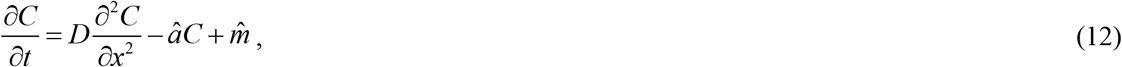

where

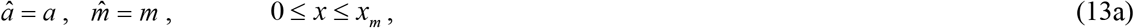

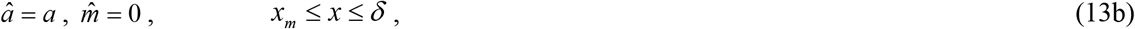

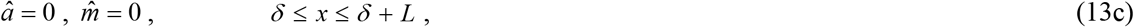

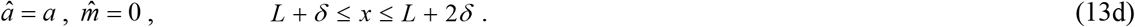

As Eq. (12) applies throughout the entire computational domain, only boundary conditions at external boundaries, *x* = 0 and *x* = *L* + 2*δ*, are necessary:

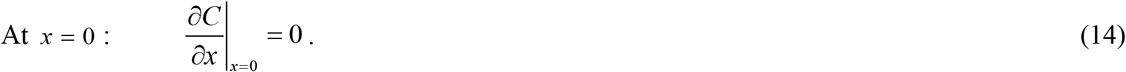

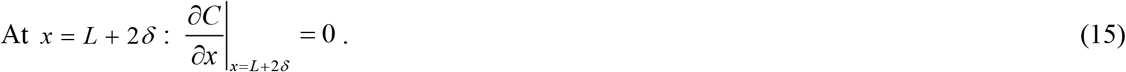

Employing the single-domain approach yields a numerical solution that automatically satisfies the continuity of the ATP concentration and flux across the interfaces between subdomains, *x* = *x*_*m*_, *x* = *δ*, and *x* = *δ* + *L*. This means that Eqs. (7)-(9) are satisfied automatically.

Eq. (12) is solved with the following initial condition:

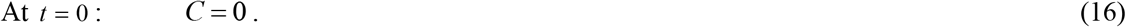

Eq. (12), along with boundary conditions given by Eqs. (14) and (15), and initial condition (16), was solved using a well-validated MATLAB PDEPE solver (MATLAB R2023a, MathWorks, Natick, MA, USA). For *f* =1 Hz, a (100,10000) mesh was employed in the (*x,t*) domain, while for *f* = 10 Hz, a (100,100000) mesh was used. At a frequency of 10 Hz, the computational time on a PC with an Intel Core i9 processor was approximately 6 hours, due to the large number of oscillations in the ATP consumption rate imposed to simulate neuronal firing.

## 3. Results

Figs. 3-5 show the results for *L*=3 μm and *f* =1 Hz. To check the accuracy of the numerical solution for ATP concentration, it was compared with the analytical solution obtained in ref. [19] for the scenario involving a constant rate of ATP consumption in a bouton (*a* = *a*_0_ = constant). For the paper to be self-contained, this solution is provided in Section S1 of the Supplementary Materials. Both the numerical and analytical solutions are identical (Fig. 3).

**Fig. 3.**
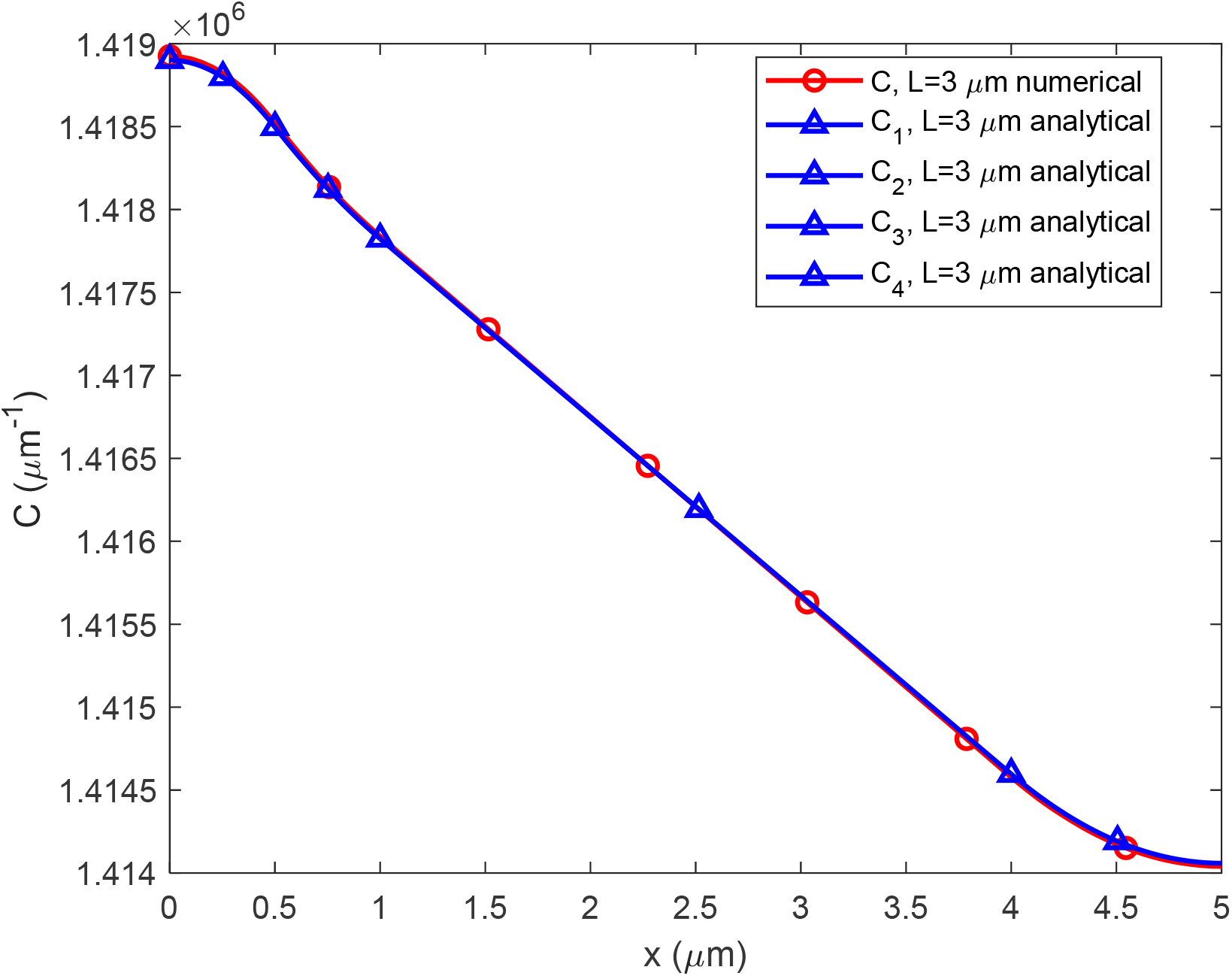
Comparison between the numerical results (this study) and the analytical solution obtained in ref. [19] for the linear ATP concentration in the axon, denoted as *C*, plotted against the distance from a stationary mitochondrion, *x*. Constant rate of ATP consumption in a bouton, *L*=3 μm.

Oscillations in the ATP consumption rate caused by neuronal firing led to small oscillations in ATP concentrations in boutons. The peak-to-peak amplitude of these oscillations is approximately 5% of their average value (Fig. 4, Table 2). The red and blue lines in Fig. 4a depict rapidly oscillating curves, resembling tight sine waves. Magnified segments of these curves, showing the last 10 seconds of the simulation, are displayed in Fig. 4b. The ATP concentration in a bouton lacking a stationary mitochondrion oscillates similarly to that in a bouton having a stationary mitochondrion, but the average concentration is approximately 0.3% lower (Fig. 4b, Table 4). However, due to the small amplitude of these oscillations, they are unable to reduce the ATP concentration in a bouton lacking a stationary mitochondrion below the threshold required to sustain synaptic activity (3.78 ×10^5^ ATP/μm, as shown in Table 1). This is explained by the high diffusivity of ATP (300 μm^2^ s^-1^, see Table 1). The oscillations depicted in Fig. 4 are consistent with the ATP concentration oscillations reported in ref. [18], which investigate the depletion of intracellular resources such as ATP due to prolonged neuronal firing and the restoration of these resources by transitioning into the down state.

**Table 2.** Effect of the average spacing between varicosities and the frequency at which neuron fires. The peak-to-peak amplitude of oscillations of the ATP concentrations because of neuron firing, percentage of the average value.

| | $L=3\ \mu\text{m}$ | $L=10\ \mu\text{m}$ |
| --- | --- | --- |
| $f = 1\ \text{Hz}$ | 5% | 2% |
| $f = 10\ \text{Hz}$ | 0.5% | 0.4% |
| $f = 200\ \text{Hz}$ | 0.06% | 0.06% |

**Fig. 4.**
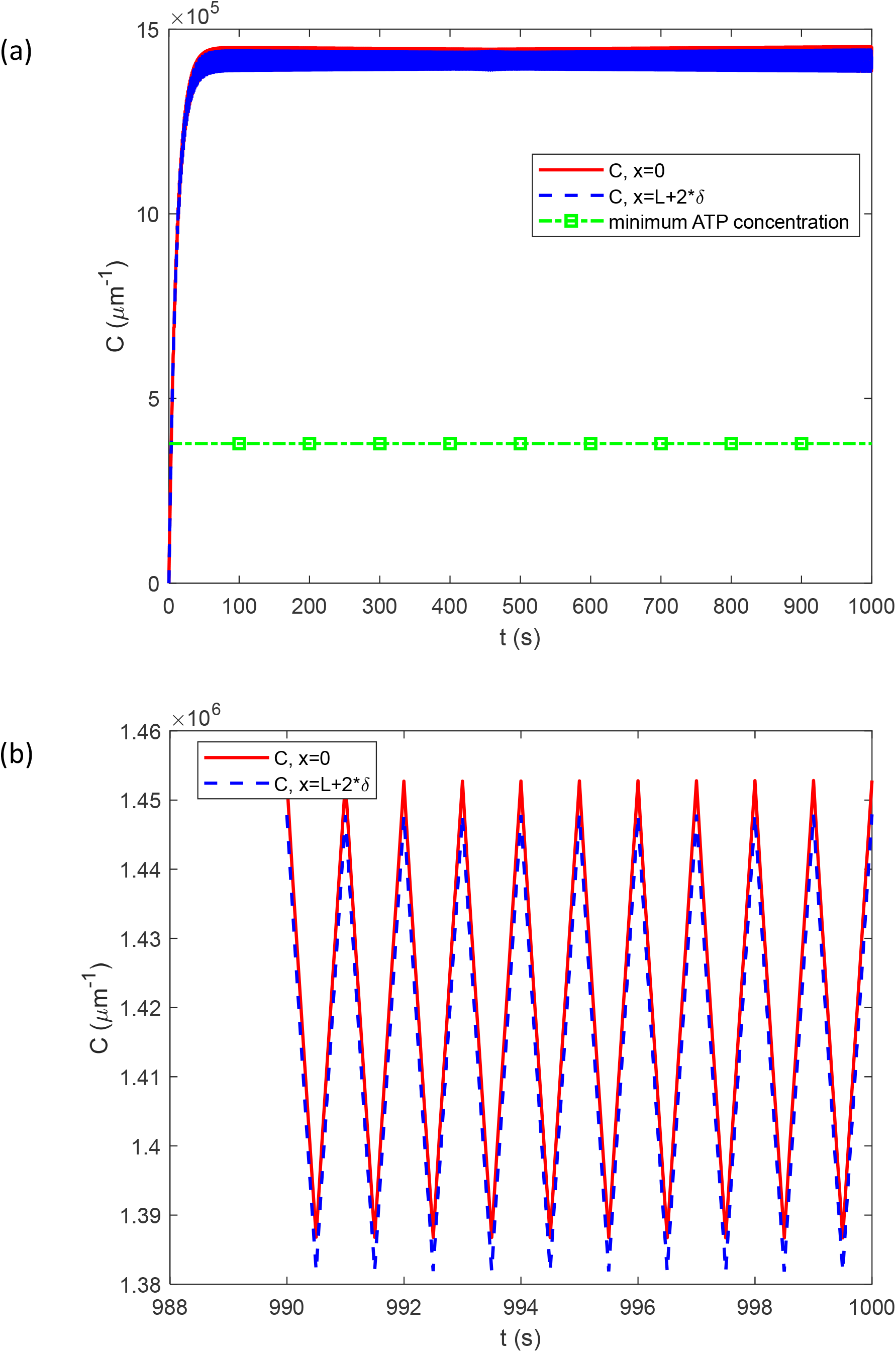
Variation with time of the linear ATP concentration at *x*=0 (at the center of a bouton with a stationary mitochondrion) and at *x* = *L* + 2*δ* (at the center of a bouton without a stationary mitochondrion). (a) The simulation covers the entire 1000 s. A line marked with squares represents the minimum linear ATP concentration required to support synaptic transmission (see Table 1). (b) The last 10 s of the simulation. *f* =1 Hz, *L*=3 μm.

The distribution for a constant ATP consumption rate is characterized by a middle curve marked with triangles. In the scenario with an oscillating ATP consumption rate, ATP concentration fluctuates with an amplitude of 2.5% around the concentration corresponding to a constant ATP consumption rate (Fig. 5, Table 3). The upper curve in Fig. 5, marked with squares, corresponds to the peak concentration time in Fig. 4b, while the lower curve in Fig. 5, marked with circles, corresponds to the minimum concentration time in Fig. 4b.

**Table 3.** Effect of the average spacing between varicosities and the frequency at which neuron fires. The amplitude, expressed as a percentage of the average value, of ATP concentration fluctuation due to oscillating ATP consumption around that observed under constant ATP consumption.

| | $L=3\ \mu\text{m}$ | $L=10\ \mu\text{m}$ |
| --- | --- | --- |
| $f = 1\ \text{Hz}$ | 2.5% | 1% |
| $f = 10\ \text{Hz}$ | 0.25% | 0.2% |
| $f = 200\ \text{Hz}$ | 0.03% | 0.03% |

**Fig. 5.**
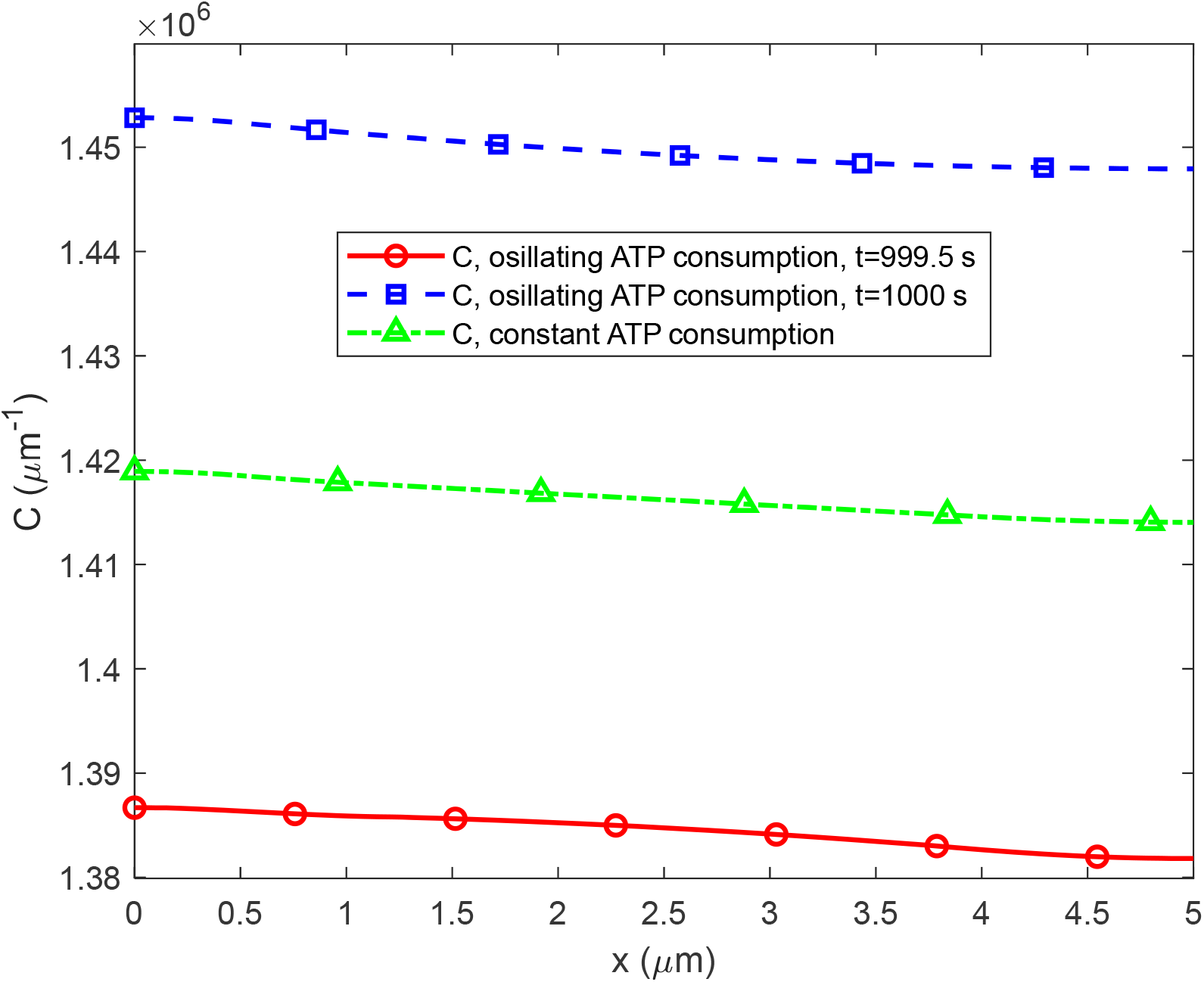
The variation of ATP concentration between the centers of two boutons, one containing a stationary mitochondrion (at *x*=0) and the other without a stationary mitochondrion (at *x* = *L* + 2*δ*), at *t*=1000 s. *f* = 1 Hz, *L*=3 μm. In the scenario with oscillating ATP consumption Eq. (5) is utilized, while the scenario with constant ATP consumption utilizes *a* = *a*_0_ (given in Table 1). In the scenario with oscillating ATP consumption, ATP concentration is depicted at two times: one corresponding to a peak ATP concentration (*t*=1000 s, see Fig. 4b), and the other corresponding to a minimum ATP concentration (*t*=999.5 s, see Fig. 4b).

In the oscillating case, ATP concentration decreases from the center of the bouton containing a stationary mitochondrion to the center of the bouton without a stationary mitochondrion (Fig. 5). The same trend is also observed in the case with constant ATP consumption. The decrease in ATP concentration displayed in Fig. 5 qualitatively agrees with the decrease in the ATP:ADP signal ratio reported in Fig. 4E of ref. [26] as the distance from a mitochondrion increases. The difference in ATP concentrations between the bouton containing a stationary mitochondrion and the bouton lacking a stationary mitochondrion is only 0.3% of the average value (Fig. 5, Table 4). This may explain why ref. [37] observed a uniform ATP/ADP ratio along an axon.

**Table 4.** Effect of the average spacing between varicosities and the frequency at which neuron fires. The percentage by which the average ATP concentration in a bouton lacking a stationary mitochondrion is lower than that in a bouton having a stationary mitochondrion.

| | $L=3\ \mu\text{m}$ | $L=10\ \mu\text{m}$ |
| --- | --- | --- |
| $f = 1\ \text{Hz}$ | 0.3% | 0.8% |
| $f = 10\ \text{Hz}$ | 0.4% | 0.6% |
| $f = 200 \text{ Hz}$ | 0.3% | 0.8% |

At *t*=999.5 s, corresponding to the lower curve in Fig. 5, the concentration at the center of the bouton without a stationary mitochondrion (at *x*=5 μm) is 3.65 times higher than the minimum ATP concentration given in Table 1. This estimate is consistent with the one given in ref. [19], where we reported a ratio of 3.75 for the scenario involving a constant ATP consumption rate.

Increasing the frequency of neuron firing from 1 Hz to 10 Hz leads to a reduction in the peak-to-peak amplitude of ATP concentration oscillations, now 0.5% of the average value, as shown in Fig. S1b in the Supplementary Materials (also see Table 2). The percentage of ATP concentration variation due to oscillating ATP consumption relative to that observed under constant ATP consumption is approximately 0.25% (Fig. S2 and Table 3). The average ATP concentration in a bouton lacking a stationary mitochondrion is approximately 0.4% lower than that in a bouton containing a stationary mitochondrion (Fig. S2 and Table 4).

Increasing neuron firing frequency to 200 Hz results in a decrease in the peak-to-peak amplitude of ATP concentration oscillations, now at 0.06% of the average value, as depicted in Fig. S3 of the Supplementary Materials (also refer to Table 2). The percentage of ATP concentration variation attributed to oscillating ATP consumption, compared to that under constant ATP consumption, is approximately 0.03% (Fig. S3 and Table 3). Additionally, the average ATP concentration in a bouton lacking a stationary mitochondrion is roughly 0.3% lower than that in a bouton containing a stationary mitochondrion (Fig. S3 and Table 4).

Given that pathological increases in the distance between mitochondria could lead to local energy failure in boutons [15], the scenario with *L*=10 μm and *f* =1 Hz was examined in Figs. 6 and 7. The peak-to-peak amplitude of ATP concentration oscillations in boutons with and without a stationary mitochondrion is approximately 2% of their average values (Fig. 6b and Table 2). This is lower than in Fig. 4b. Increasing the distance between the boutons appears to decrease the amplitude of ATP concentration oscillations.

**Fig. 6.**
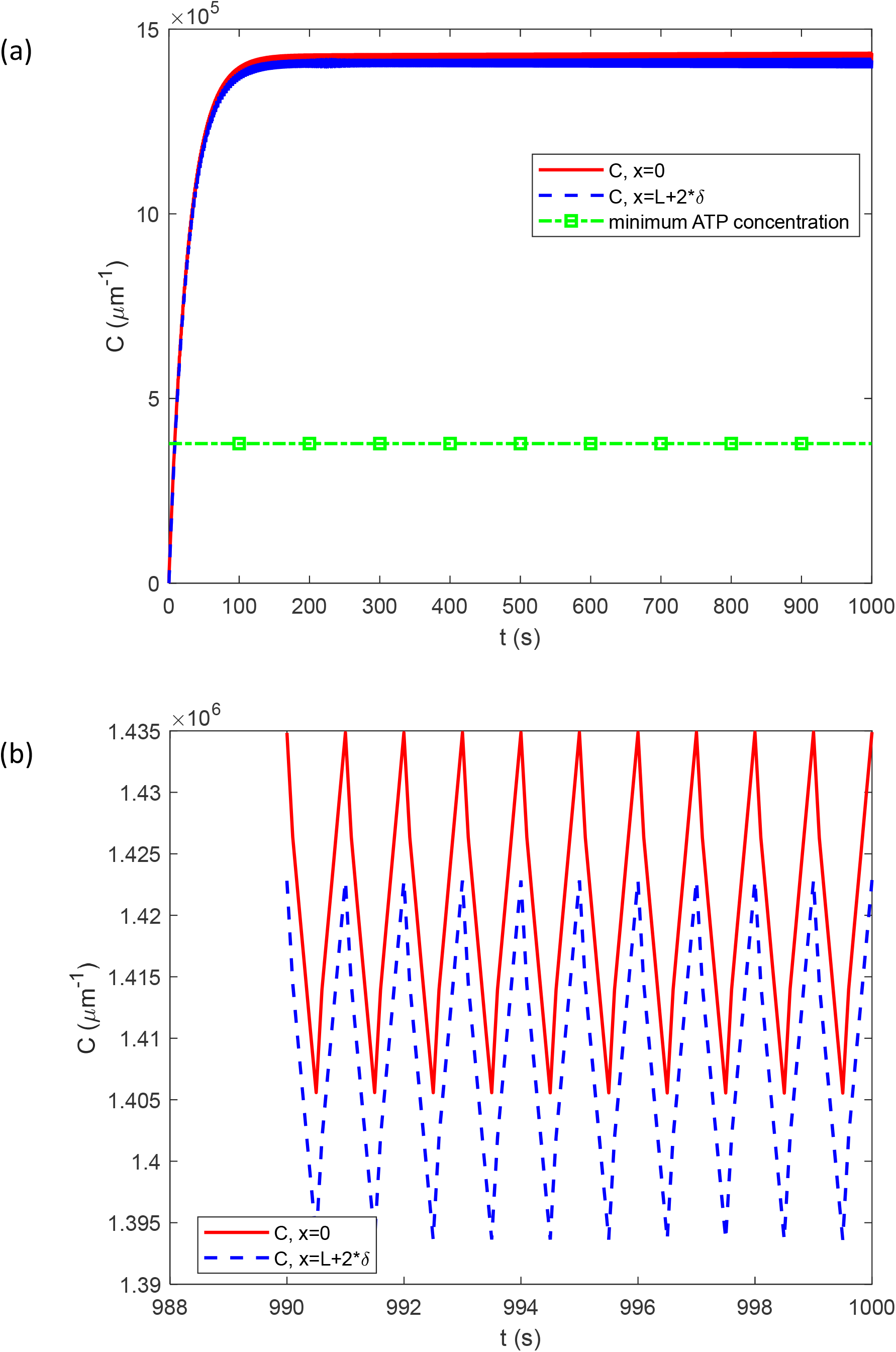
Variation with time of the linear ATP concentration at *x*=0 (at the center of a bouton with a stationary mitochondrion) and at *x* = *L* + 2*δ* (at the center of a bouton without a stationary mitochondrion). (a) The simulation covers the entire 1000 s. A line marked with squares represents the minimum linear ATP concentration required to support synaptic transmission (see Table 1). (b) The last 10 s of the simulation. *f* =1 Hz, *L*=10 μm.

**Fig. 7.**
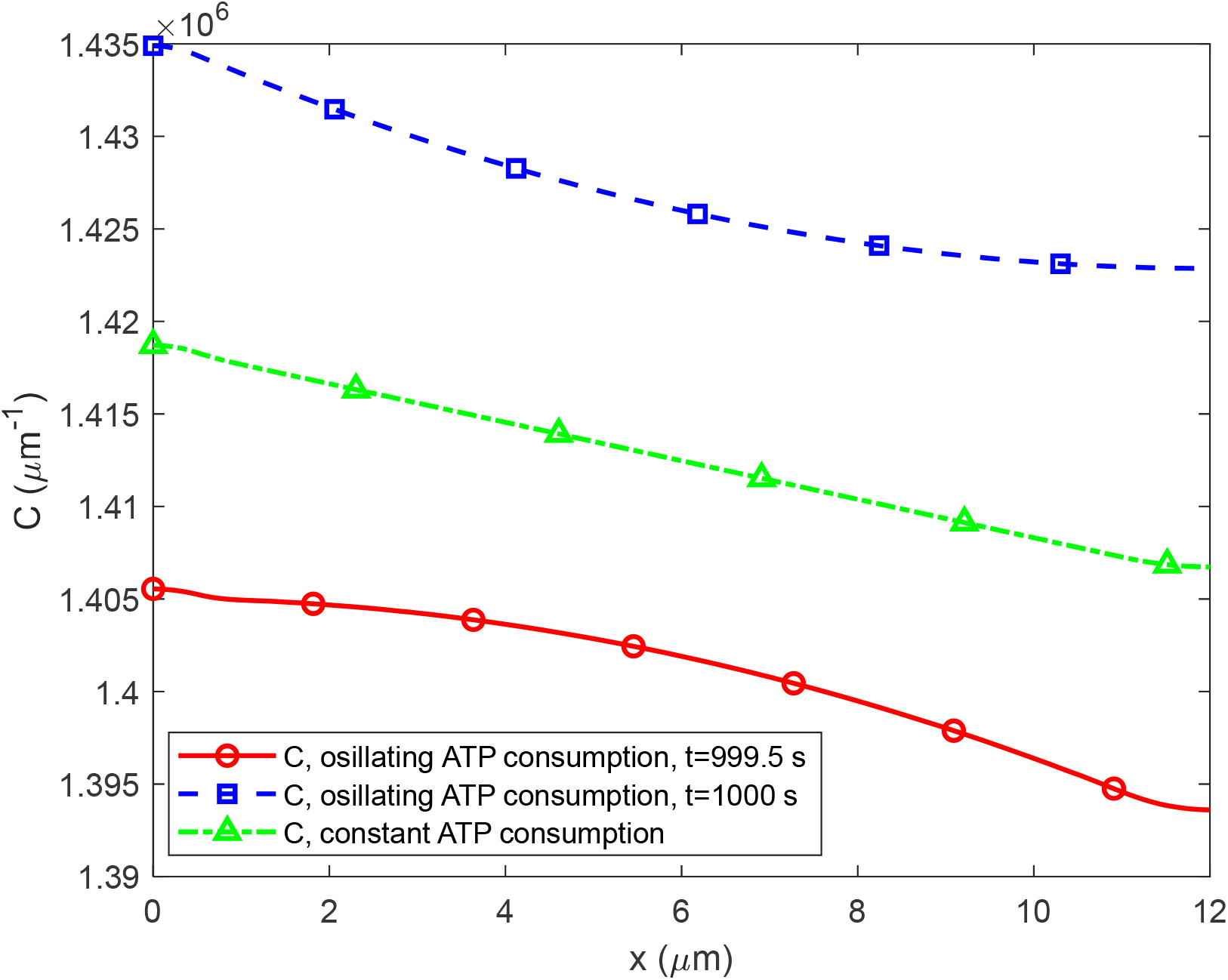
The variation of ATP concentration between the centers of two boutons, one containing a stationary mitochondrion (at *x*=0) and the other without a stationary mitochondrion (at *x* = *L* + 2*δ*), at *t*=1000 s. *f* = 1 Hz, *L*=10 μm. In the scenario with oscillating ATP consumption Eq. (5) is utilized, while the scenario with constant ATP consumption utilizes *a* = *a*_0_ (given in Table 1). In the scenario with oscillating ATP consumption, ATP concentration is shown at two times: one corresponding to a peak ATP concentration (*t*=1000 s, see Fig. 6b), and the other corresponding to a minimum ATP concentration (*t*=999.5 s, see Fig. 6b).

For *L*=10 μm and *f* =1 Hz, in the case of an oscillating ATP consumption rate, ATP concentration fluctuates with an amplitude of approximately 1% around the curve corresponding to constant ATP consumption rate (Fig. 7 and Table 3). It is noteworthy that this amplitude is smaller compared to Fig. 5, where it was 2.5% (see Table 3), suggesting that increasing the distance between the boutons decreases the amplitude of ATP concentration variation.

In the oscillating scenario, at a given time, the ATP concentration decreases from the center of the bouton with a stationary mitochondrion to the one without (Fig. 7). The difference in ATP concentration between a bouton containing a stationary mitochondrion and a bouton lacking a stationary mitochondrion is 0.8% (Fig. 7 and Table 4). The increased difference in ATP concentrations between boutons with and without a stationary mitochondrion (previously 0.3% in Fig. 5, also see Table 4) is attributed to diffusion-driven transport of ATPs over a greater distance. For the lower curve in Fig. 7, the ATP concentration in a bouton lacking a stationary mitochondrion remains 3.69 times higher than the minimum ATP concentration required to sustain synaptic activity (3.78 ×10^5^ ATP/μm).

Increasing the frequency of neuron firing from 1 Hz to 10 Hz results in a reduced peak-to-peak amplitude of ATP concentration oscillations (0.4%, as shown in Fig. S4b, see also Table 2). When considering an oscillating ATP consumption rate, the amplitude of ATP concentration variation around the curve corresponding to a constant consumption rate ranges approximately from 0.2% to 0.25% (Fig. S5 and Table 3). The lack of symmetry between the upper and lower curves, corresponding to the minimum and maximum concentrations for the oscillating case in Fig. S5, occurs because the minimum concentration for the oscillating case does not occur precisely at 999.95 s. At a given time, the ATP concentration in a bouton lacking a stationary mitochondrion is between 0.4% to 0.6% lower than that in a bouton containing one, depending on the distance between the boutons (Fig. S5, also see Table 4).

For *L*=10 μm and *f* = 200 Hz, the peak-to-peak amplitude of ATP concentration oscillations decreases compared to the case with *f* =10 Hz, now at 0.06% of the average value, as depicted in Fig. S6 of the Supplementary Materials (also refer to Table 2). The percentage of ATP concentration variation attributed to oscillating ATP consumption, compared to that under constant ATP consumption, is approximately 0.03% (Fig. S6 and Table 3). Additionally, the average ATP concentration in a bouton lacking a stationary mitochondrion is roughly 0.8% lower than that in a bouton containing a stationary mitochondrion (Fig. S6 and Table 4).

The increase in ATP diffusivity, *D*, by a factor of 10 did not alter the amplitude of ATP concentration oscillations, which remained at 5% for *f* = 1 Hz (Table S1). However, differences in ATP concentration between boutons with and without stationary mitochondria became negligible, as shown by horizontal lines in Fig. 8, attributed to the increased ATP diffusivity.

**Fig. 8.**
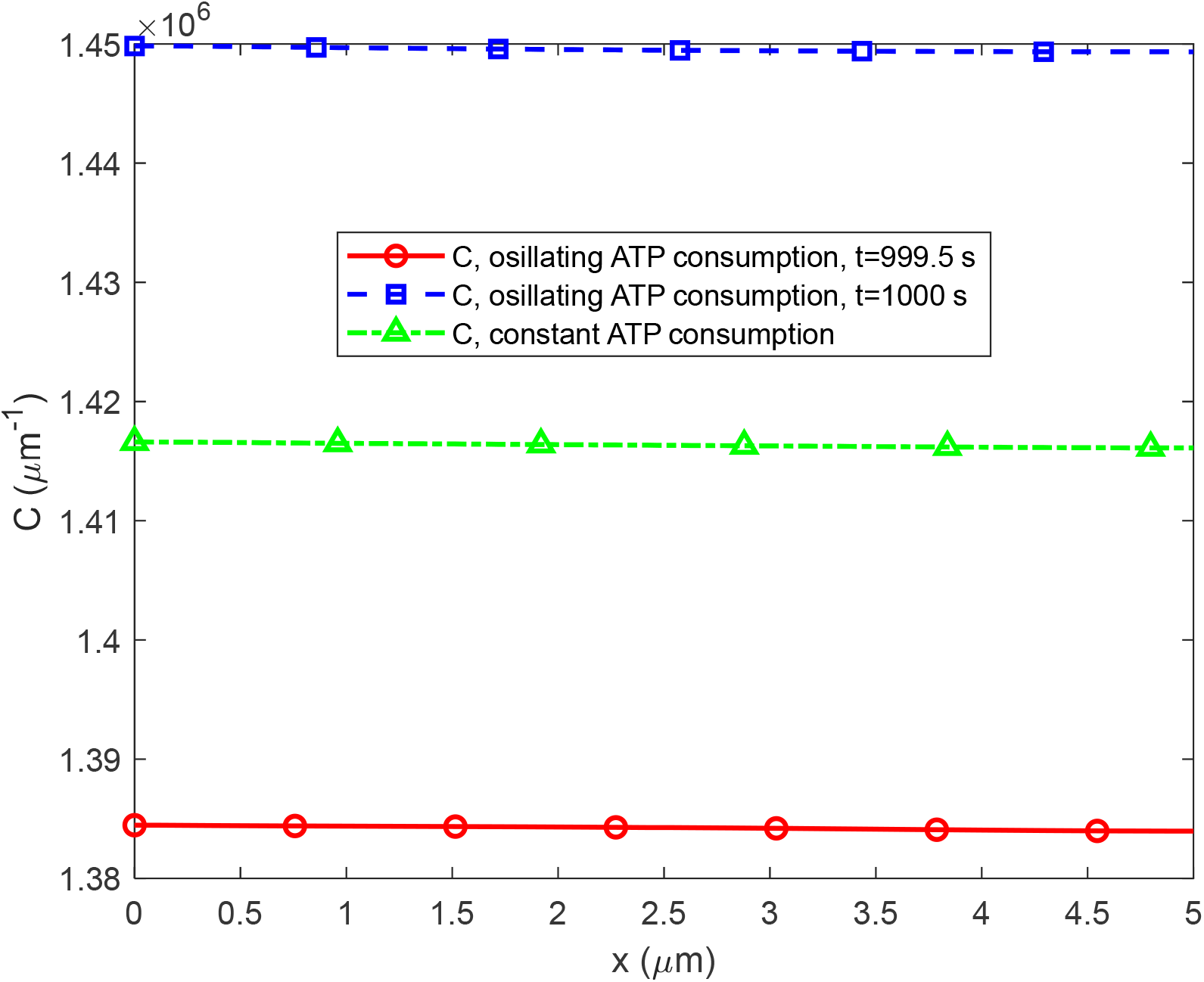
Similar to Fig. 5, but now with *f* = 1 Hz, *L*=3 μm, *D* = 3000 μm^2^ s^-1^, the values of other parameters are as listed in Table 1.

Increasing the rate of ATP production per unit length of a mitochondrion, *m*, by a factor of 10 did not alter the amplitude of ATP concentration oscillation. These oscillations remained at 5% for *f* = 1 Hz (Table S2). However, for *f* =1 Hz this modification increased ATP concentration by an order of magnitude (Fig. 9).

**Fig. 9.**
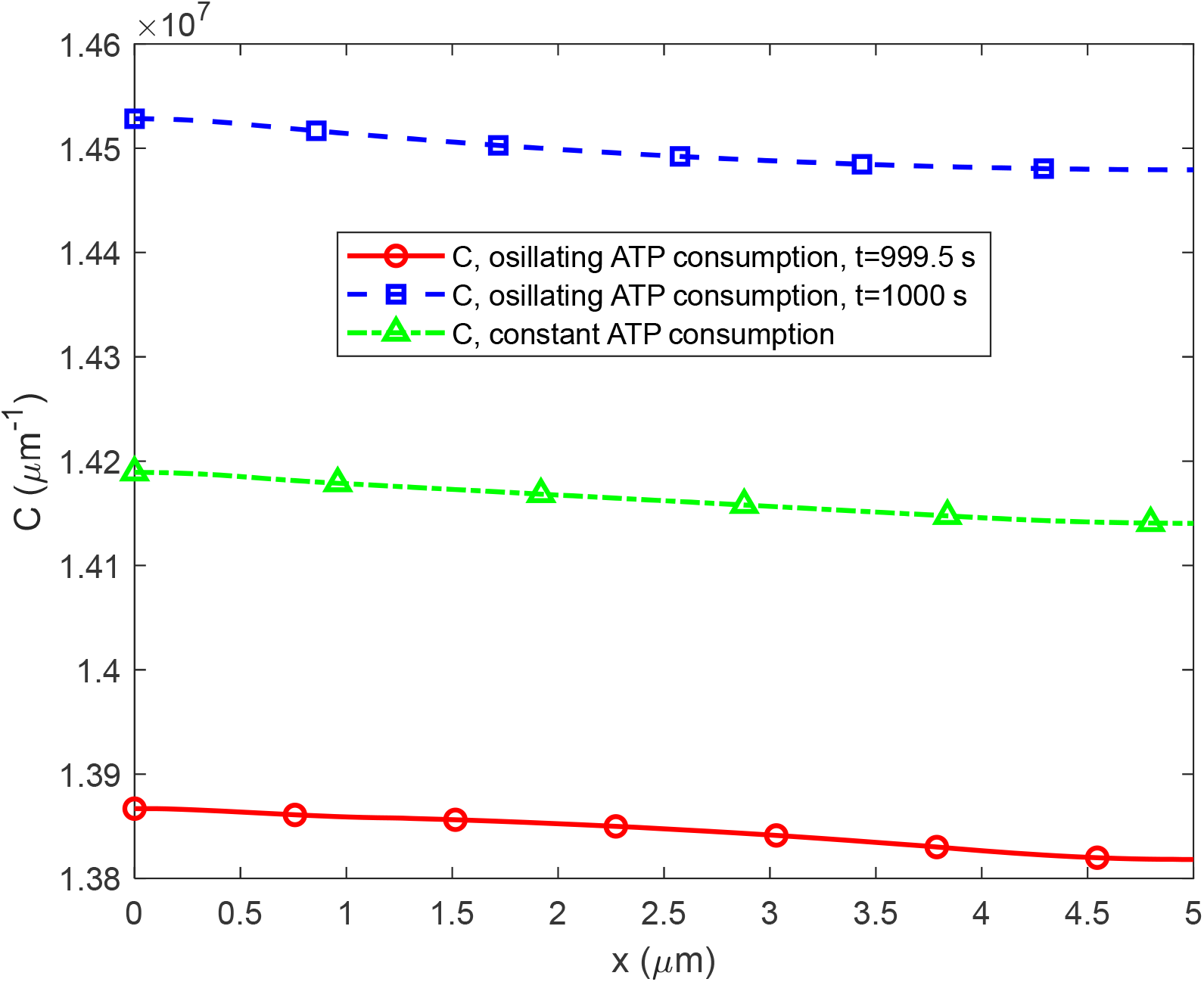
Similar to Fig. 5, but now with *f* = 1 Hz, *L*=3 μm, *m*=1.29 ×10^7^ μm^-1^s^-1^, the values of other parameters are as listed in Table 1.

The increase in the kinetic constant describing the rate of ATP consumption in boutons, *a*_0_, by a factor of 10 increased the amplitude of ATP concentration oscillations (calculated as a percentage of the average value) from 5% to 44% for *f* = 1 Hz (Table S3). This modification also decreased ATP concentration by a factor of 8 for *f* =1 Hz, as depicted in Fig. 10, which is attributed to larger consumption of ATP in boutons.

**Fig. 10.**
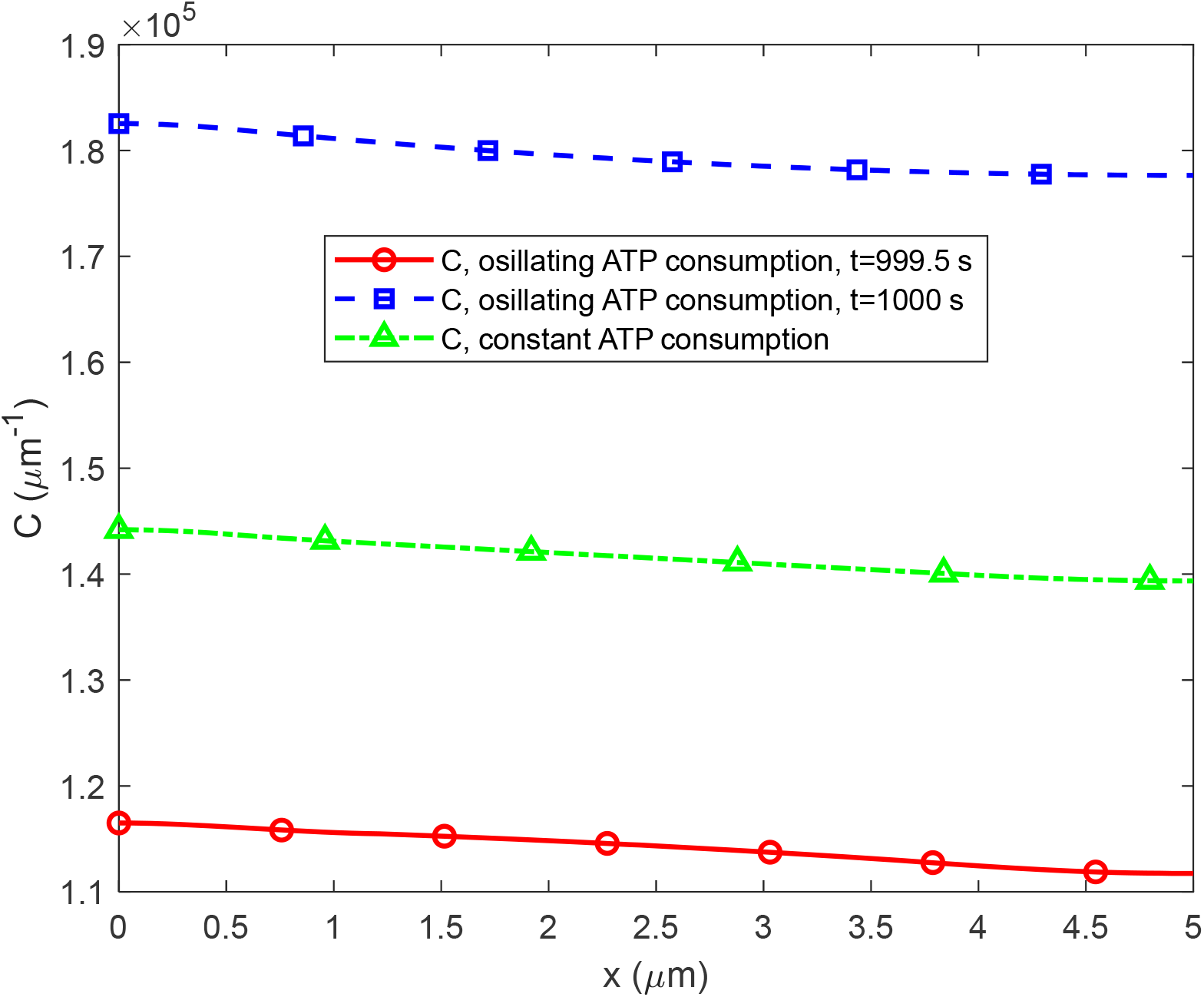
Similar to Fig. 5, but now with *f* = 1 Hz, *L*=3 μm, *a*_0_ =2.3 s^-1^, the values of other parameters are as listed in Table 1.

It should be noted that increasing the parameters *D, m*, and *a*_0_ by a factor of 10 cannot be considered physiologically relevant; this test was performed only to investigate the sensitivity of the model to these parameters.

## 4. Discussion, limitations of the model, and future directions

The study examined ATP concentration oscillations in boutons triggered by neuron firing. The peak-to-peak amplitudes of these oscillations ranged from 0.06% to 5% of their average values, depending on the intervaricosity spacing and the frequency of neuron firing (Table 2). At low oscillation frequencies (1-10 Hz), increasing the intervaricosity spacing and the frequency of neuron firing reduces the amplitude of ATP concentration oscillations. However, at large oscillation frequencies (200 Hz), intervaricosity spacing does not affect the amplitude of ATP concentration oscillations. The oscillations are observed to be similar in boutons containing and lacking a stationary mitochondrion.

When considering an oscillating ATP consumption rate, the ATP concentration fluctuates about that computed assuming a constant ATP consumption rate with an amplitude between 0.03% and 2.5% of the value corresponding to a constant ATP consumption rate (Table 3). At low oscillation frequencies (1-10 Hz), increasing the spacing between varicosities and the frequency of neuron firing reduces the disparity in ATP concentrations between scenarios with oscillating and constant ATP consumption rates. However, at high oscillation frequencies (200 Hz), the spacing between varicosities does not affect the difference in ATP concentrations between scenarios with oscillating and constant ATP consumption rates.

Diffusion-driven transport necessitates concentration gradients, requiring an ATP gradient between boutons with and without a stationary mitochondrion. Results indicate between a 0.3% to a 0.8% lower ATP concentration in boutons lacking stationary mitochondria (Table 4). The increase in intervaricosity spacing increases the difference in ATP concentrations between boutons containing and lacking a stationary mitochondrion. However, the ATP concentration in these boutons remains approximately 3.75 times higher than that needed for synaptic activity, suggesting that having a stationary mitochondrion in every bouton is not necessary. With a mitochondrion present in every second bouton, ATP diffusion maintains sufficient ATP concentration levels, avoiding significant concentration drops in boutons lacking mitochondria. The frequency of neuron firing does not significantly affect the difference in ATP concentrations between boutons containing and lacking a stationary mitochondrion. This difference is primarily determined by ATP diffusion, which remains unaffected by the oscillations in ATP demand.

The main effect of increasing the ATP diffusivity is the reduction in the difference in ATP concentration between a bouton containing a stationary mitochondrion and a bouton lacking a stationary mitochondrion. The main effect of increasing the rate of ATP production per unit length of a mitochondrion is an increase in ATP concentration. Increasing the kinetic constant describing the rate of ATP consumption in boutons decreases the ATP concentration and increases the amplitude of ATP concentration oscillations relative to the average value.

The model’s limitations stem from assuming a constant rate of ATP production by mitochondria and neglecting ATP production by glycolysis. Future research should aim to develop more comprehensive models of biochemical processes within synapses and mitochondria to better simulate ATP production and consumption rates.

## Abbreviations

CV: control volume
PDE: partial differential equation

## Acknowledgment

AVK acknowledges the support provided by the National Science Foundation (grant CBET-2042834) and the Alexander von Humboldt Foundation through the Humboldt Research Award.

## Supplementary Materials

### S1. Analytical solutions of governing equations obtained in ref. [19]

The analytical solution of Eqs. (1)-(4) with boundary conditions (6)-(10) for a constant rate of energy consumption in boutons (*a* = *a*_0_) at steady-state is given by the following equations:

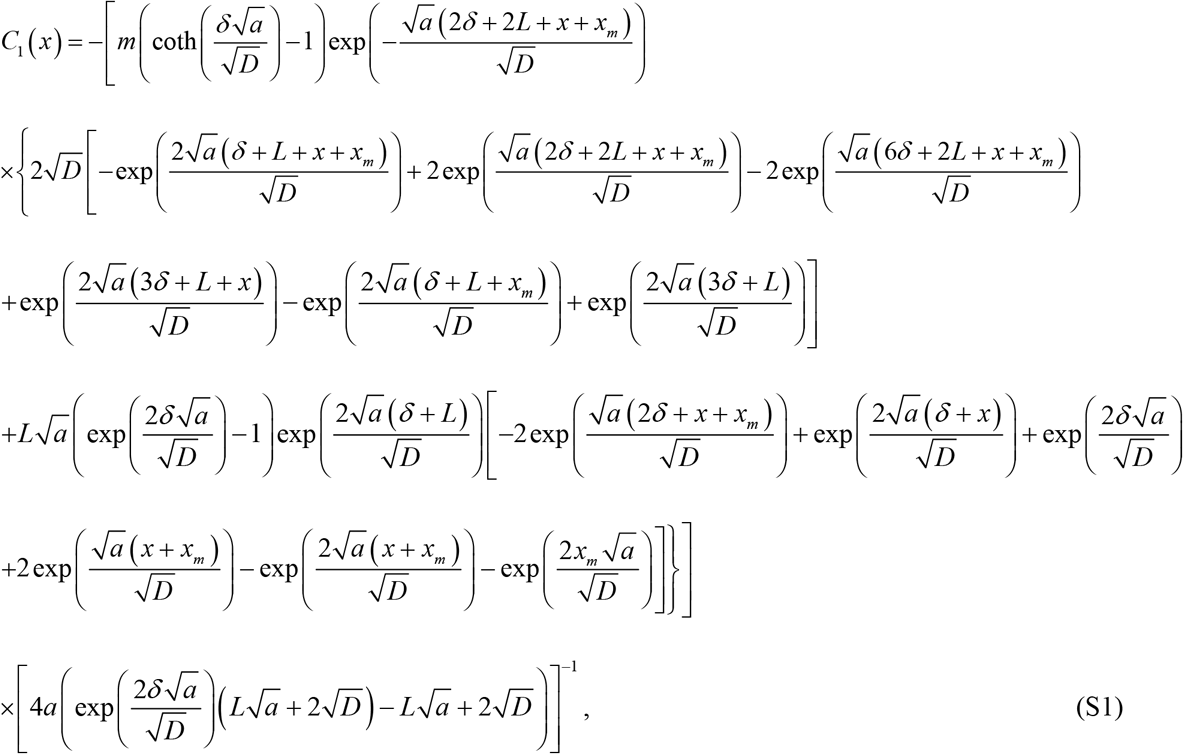

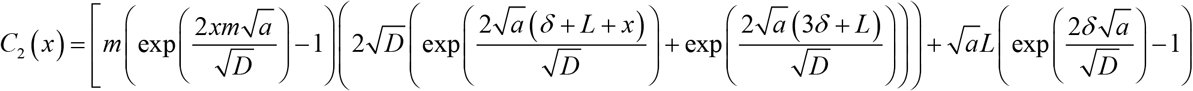

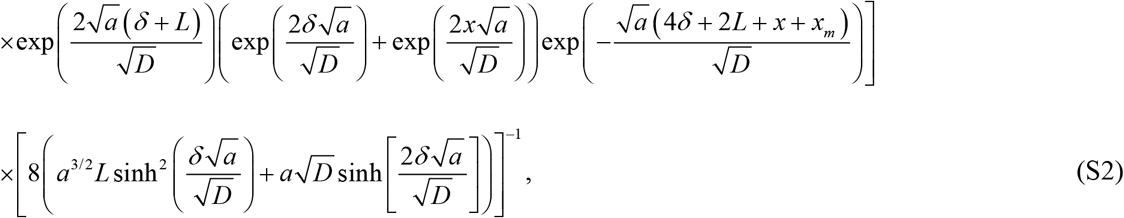

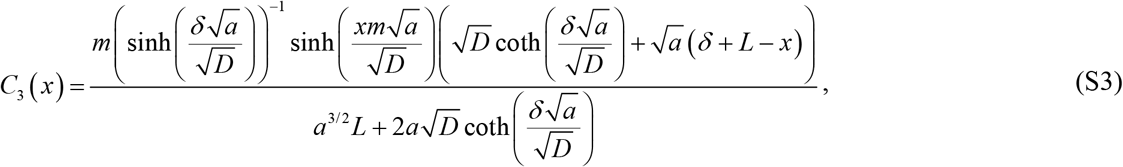

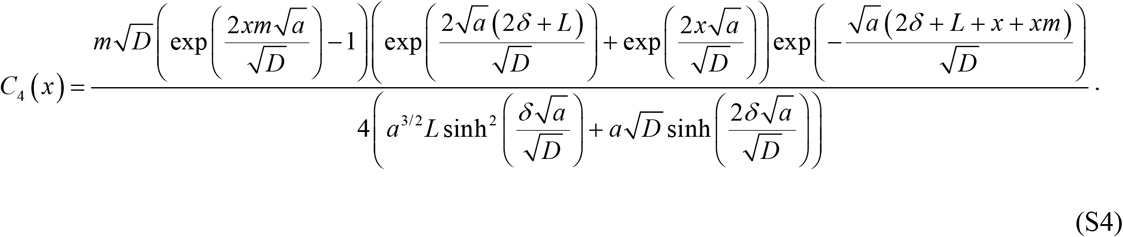

### S2. Supplementary tables

**Table S1.** Effect of the diffusivity of ATP in the cytosol. The peak-to-peak amplitude of oscillations of the ATP concentrations because of neuron firing, percentage of the average value.

| | $D=300 \mu\text{m}^2 \text{s}^{-1}$ | $D=3000 \mu\text{m}^2 \text{s}^{-1}$ |
| --- | --- | --- |
| $f = 1 \text{ Hz}$ | 5% | 5% |

**Table S2.** Effect of the rate of ATP production per unit length of a mitochondrion. The peak-to-peak amplitude of oscillations of the ATP concentrations because of neuron firing, percentage of the average value.

| | $m=1.29 \times 10^6 \mu\text{m}^{-1} \text{s}^{-1}$ | $m=1.29 \times 10^7 \mu\text{m}^{-1} \text{s}^{-1}$ |
| --- | --- | --- |
| $f = 1 \text{ Hz}$ | 5% | 5% |

**Table S3.**
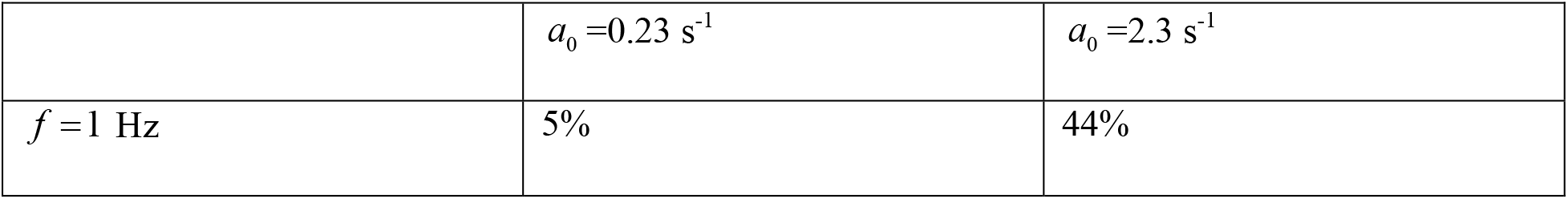
Effect of the kinetic constant describing the rate of ATP consumption in a bouton. The peak-to-peak amplitude of oscillations of the ATP concentrations because of neuron firing, percentage of the average value.

| | $a_0 = 0.23 \text{ s}^{-1}$ | $a_0 = 2.3 \text{ s}^{-1}$ |
| --- | --- | --- |
| $f = 1 \text{ Hz}$ | 5% | 44% |

### S3. Supplementary figures

**Fig. S1.**
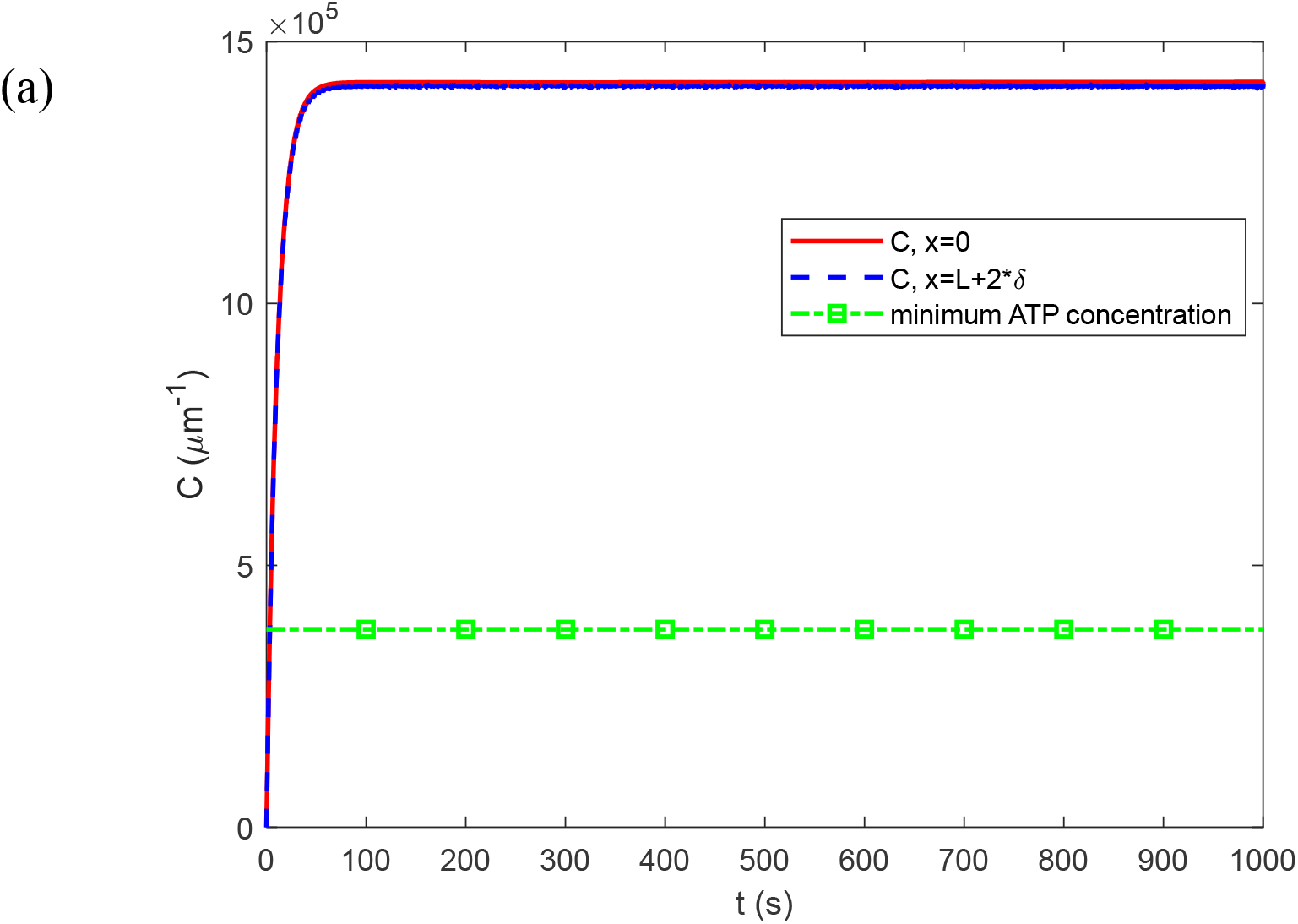
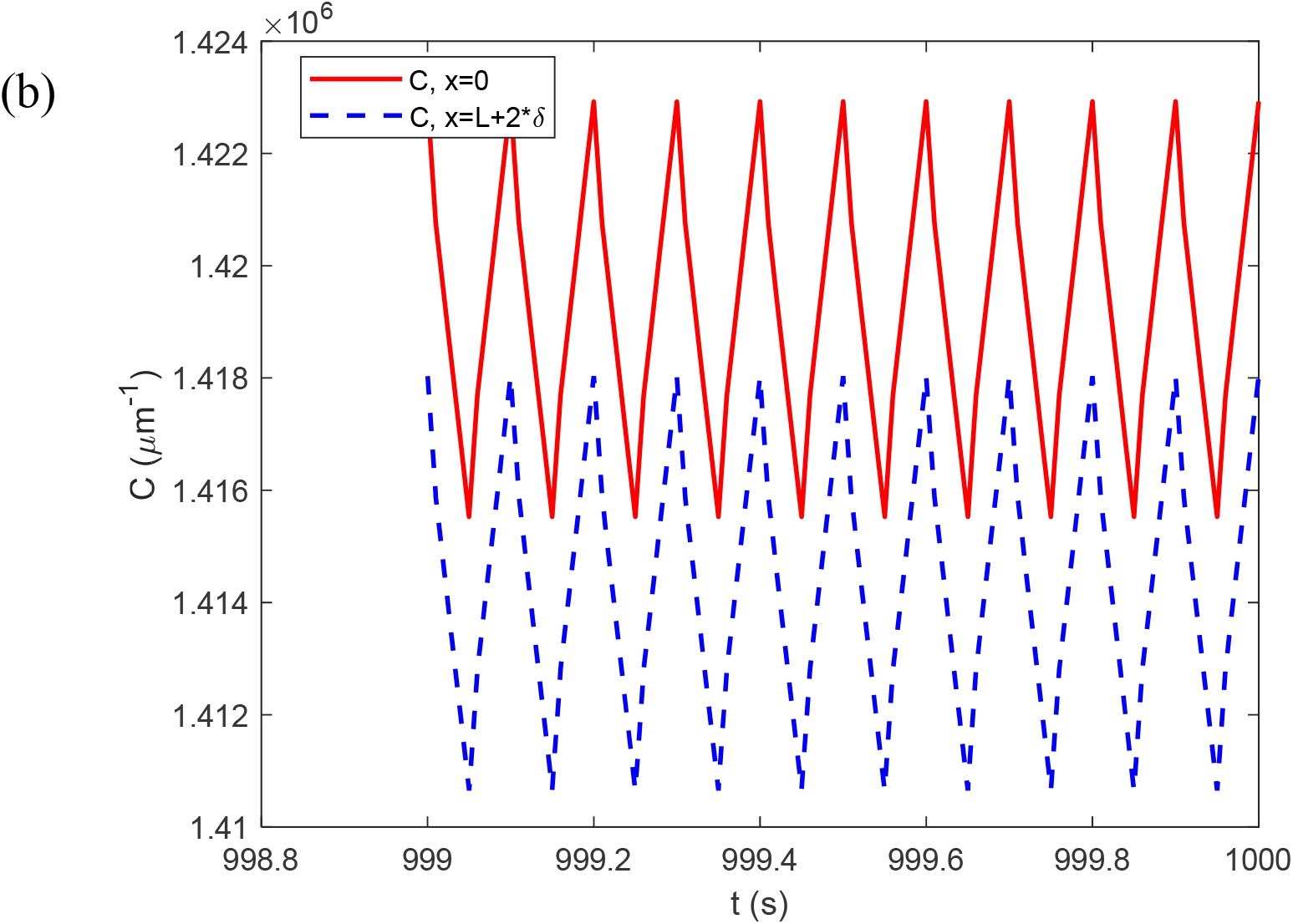
Variation with time of the linear ATP concentration at *x*=0 (at the center of a bouton with a stationary mitochondrion) and at *x* = *L* + 2*δ* (at the center of a bouton without a stationary mitochondrion). (a) The simulation covers the entire 1000 s. The line marked with squares represents the minimum linear ATP concentration required to support synaptic transmission (see Table 1). (b) The last 1 s of the simulation. *f* = 10 Hz, *L*=3 μm.

**Fig. S2.**
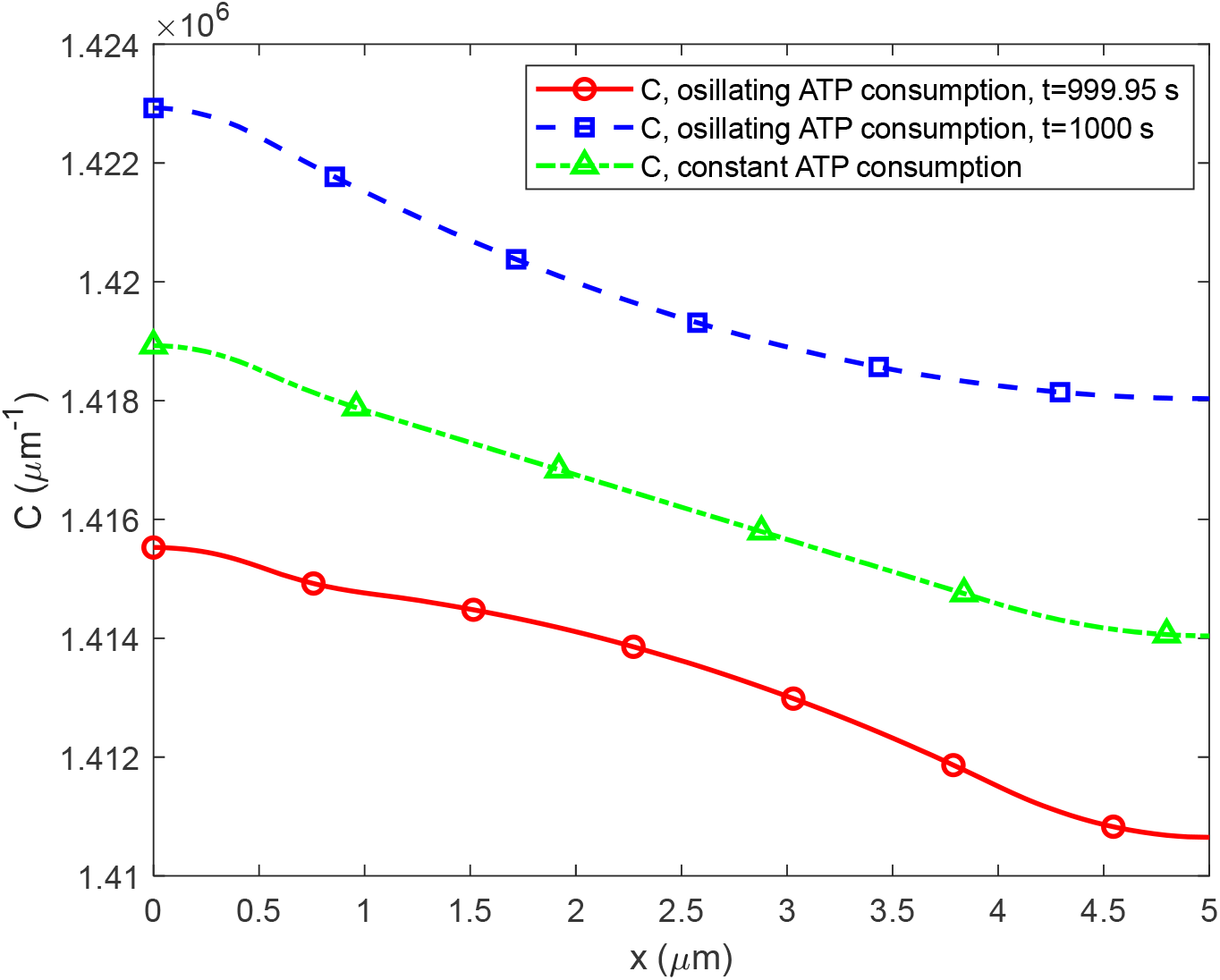
The variation of ATP concentration between the centers of two boutons, one containing a stationary mitochondrion (at *x*=0) and the other without a stationary mitochondrion (at *x* = *L* + 2*δ*), at *t*=1000 s. *f* = 10 Hz, *L*=3 μm. In the scenario with oscillating ATP consumption Eq. (5) is utilized, while the scenario with constant ATP consumption utilizes *a* = *a*_0_ (given in Table 1). In the scenario with oscillating ATP consumption, ATP concentration is shown at two times: one corresponding to a peak ATP concentration (*t*=1000 s, see Fig. S1b), and the other corresponding to a minimum ATP concentration (*t*=999.95 s, see Fig. S1b).

**Fig. S3.**
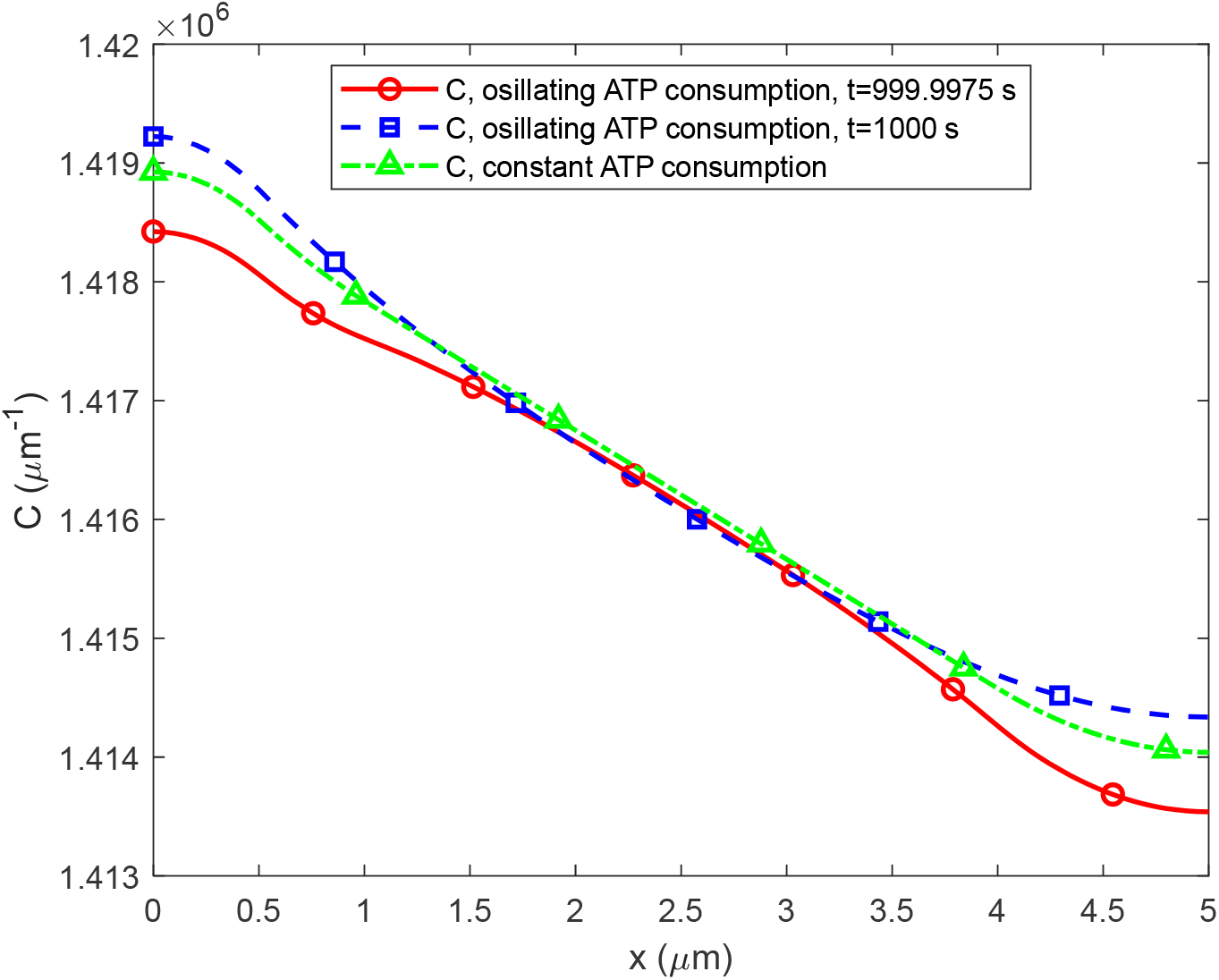
The variation of ATP concentration between the centers of two boutons, one containing a stationary mitochondrion (at *x*=0) and the other without a stationary mitochondrion (at *x* = *L* + 2*δ*), at *t*=1000 s. *f* = 200 Hz, *L*=3 μm. In the scenario with oscillating ATP consumption Eq. (5) is utilized, while the scenario with constant ATP consumption utilizes *a* = *a*_0_ (given in Table 1). In the scenario with oscillating ATP consumption, ATP concentration is shown at two times: one corresponding to a maximum ATP concentration (*t*=1000 s), and the other corresponding to the closest minimum ATP concentration (*t*=999.9975 s).

**Fig. S4.**
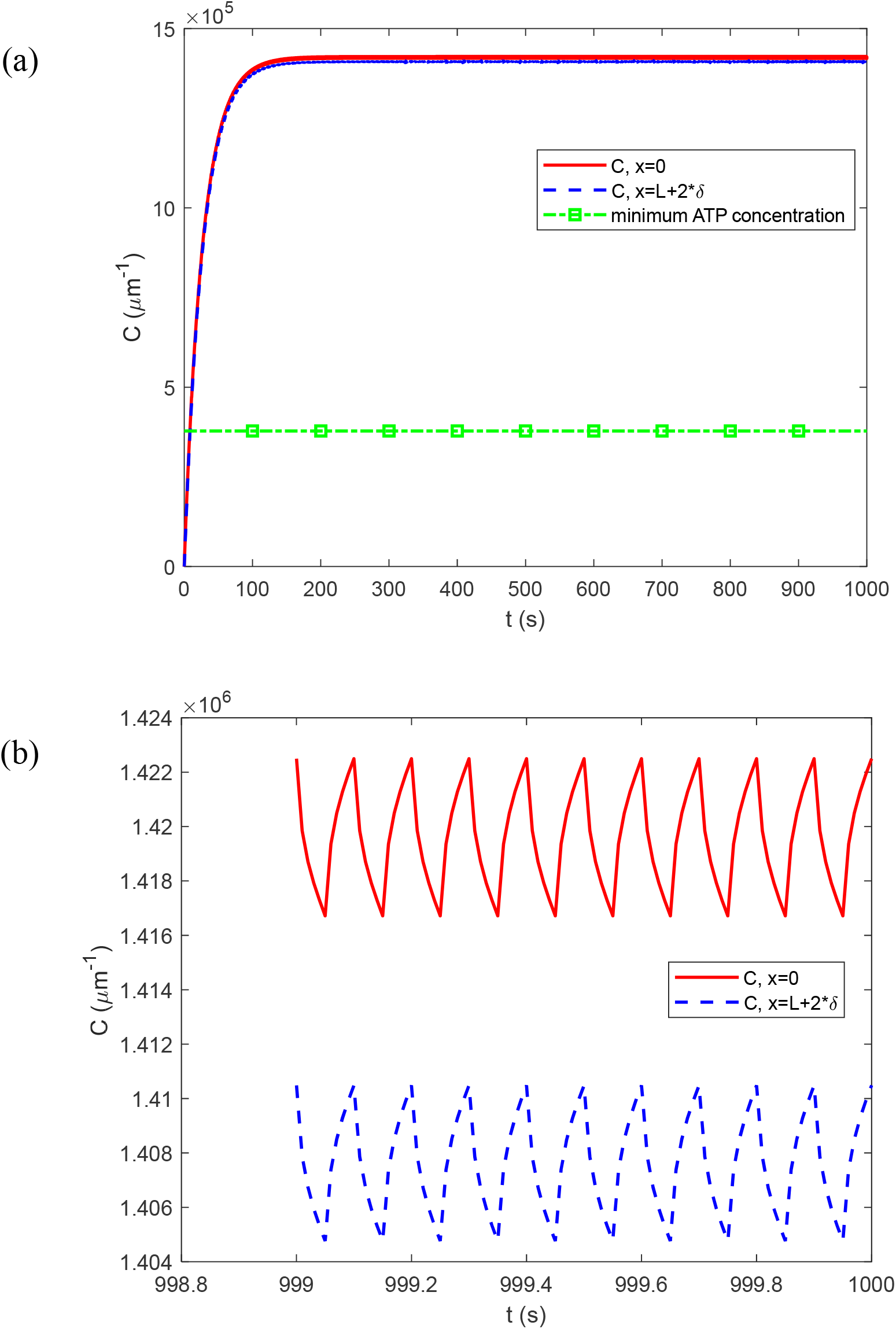
Variation with time of the linear ATP concentration at *x*=0 (at the center of a bouton with a stationary mitochondrion) and at *x* = *L* + 2*δ* (at the center of a bouton without a stationary mitochondrion). (a) The simulation covers the entire 1000 s. The line marked with squares represents the minimum linear ATP concentration required to support synaptic transmission (see Table 1). (b) The last 1 s of the simulation. *f* = 10 Hz, *L*=10 μm.

**Fig. S5.**
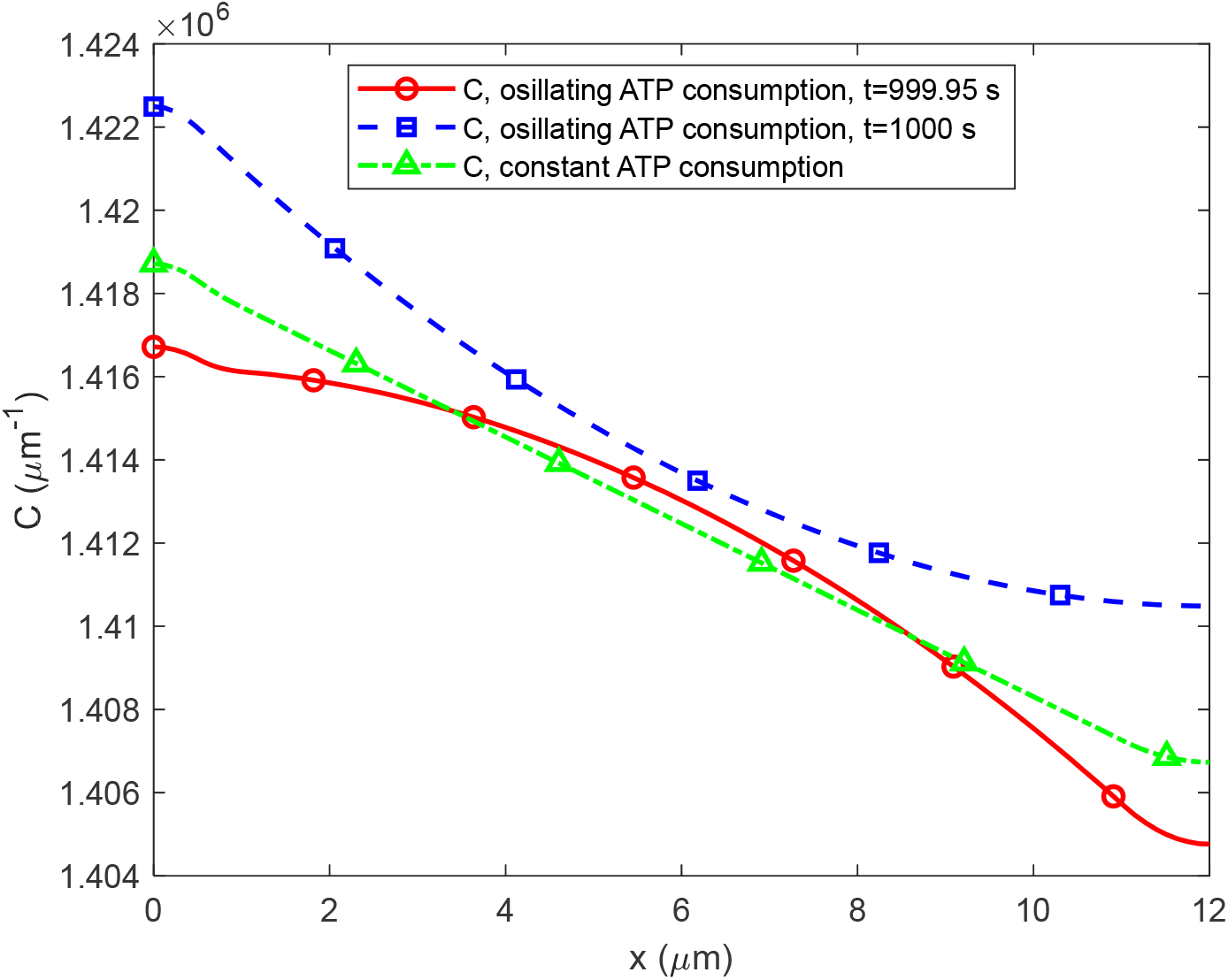
The variation of ATP concentration between the centers of two boutons, one containing a stationary mitochondrion (at *x*=0) and the other without a stationary mitochondrion (at *x* = *L* + 2*δ*), at *t*=1000 s. *f* = 10 Hz, *L*=10 μm. In the scenario with oscillating ATP consumption Eq. (5) is utilized, while the scenario with constant ATP consumption utilizes *a* = *a*_0_ (given in Table 1). In the scenario with oscillating ATP consumption, the ATP concentration is shown at two times: one corresponding to a peak ATP concentration (*t*=1000 s, see Fig. S4b), and the other corresponding to a minimum ATP concentration (*t*=999.95 s, see Fig. S4b).

**Fig. S6.**
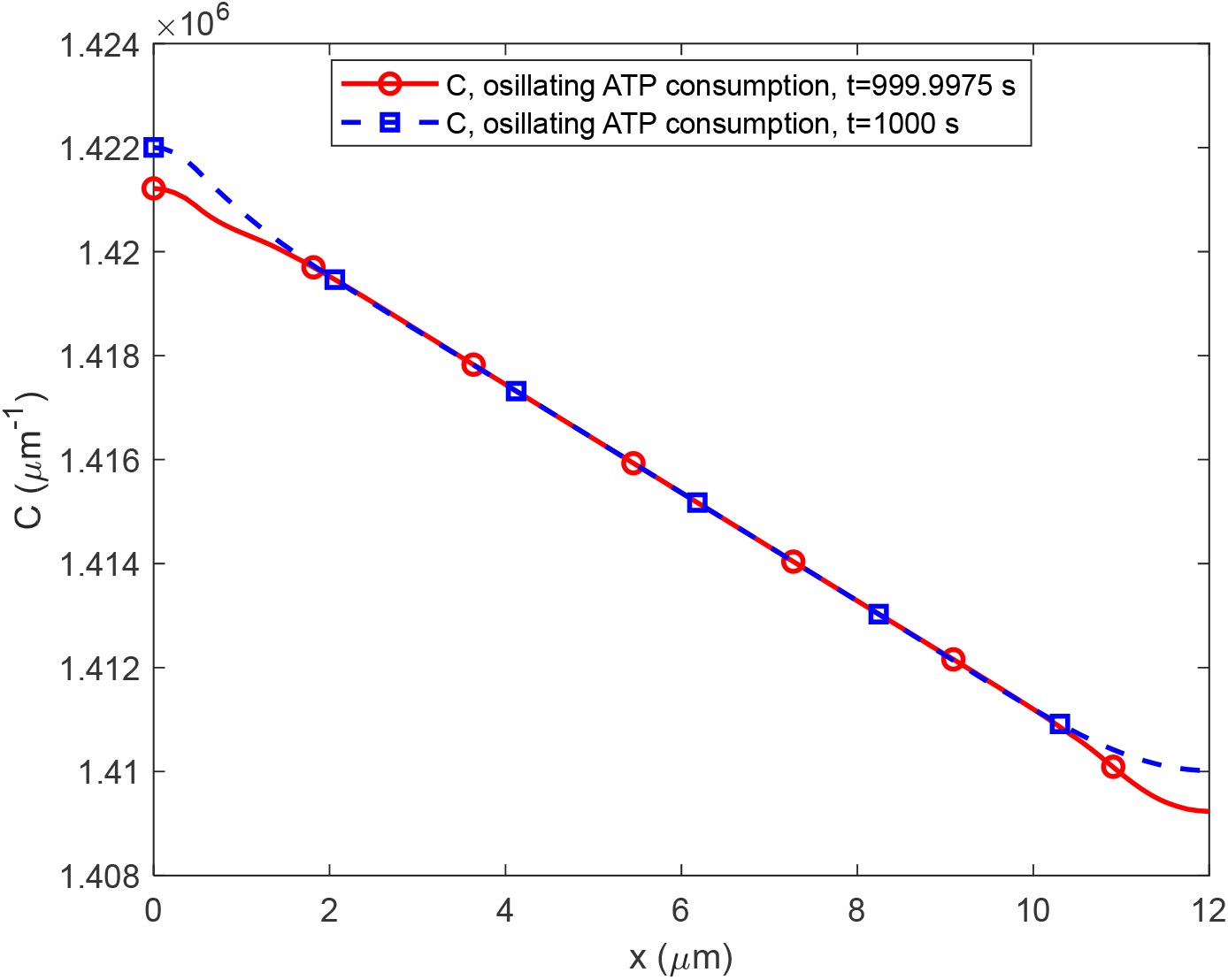
The variation of ATP concentration between the centers of two boutons, one containing a stationary mitochondrion (at *x*=0) and the other without a stationary mitochondrion (at *x* = *L* + 2*δ*), at *t*=1000 s. *f* = 200 Hz, *L*=10 μm. The scenario with oscillating ATP consumption is displayed, and Eq. (5) is employed to simulate the periodic ATP consumption rate in this scenario. ATP concentration is shown at two times: one corresponding to a maximum ATP concentration (*t*=1000 s), and the other corresponding to the closest minimum ATP concentration (*t*=999.9975 s).

